# Features fusion or not: harnessing multiple pathological foundation models using Meta-Encoder for downstream tasks fine-tuning

**DOI:** 10.1101/2025.06.05.657960

**Authors:** Ruitian Gao, Zhaochang Yang, Xin Yuan, Yifan Wang, Yujia Xia, Yufei Zhang, Bingyan Zheng, Yuqiao Gong, Yiping Yue, Zhangsheng Yu

**Author notes:** Corresponding authors: Zhangsheng Yu. These authors contributed equally to this work (Ruitian Gao, Zhaochang Yang).

## Abstract

The emergence of diverse pathological foundation models has empowered computational pathology tasks, including tumor classification, biomarker prediction, and RNA expression prediction. However, variations in model architecture and data sources lead to inconsistent downstream performance and complicate centralized training. Specifically, the lack of data sharing makes retraining foundation models with pooled data infeasible. Alternatively, the release of model parameters enables combining multiple models during fine-tuning. Inspired by the meta-analysis method, we propose the Meta-Encoder framework, which integrates features from multiple foundation models to generate a comprehensive representation, improving downstream fine-tuning task performance. Comparative experiments demonstrate that Meta-Encoder is more effective than individual foundation models, with its strengths more pronounced in handling complex tasks. While single models may perform sufficiently well for simple tasks, Meta-Encoder can match or even surpass the best-performing single model, alleviating concerns over model selection. Moderately challenging tasks benefit from Meta-Encoder’s concatenation or self-attention strategies, with the latter demonstrating superior performance in more challenging scenarios. For highly complex tasks, such as high-dimensional gene expression prediction, self-attention proves to be the most effective Meta-Encoder strategy, balancing feature integration and computational efficiency. For three patch-level spatial gene expression prediction tasks (HEST-Benchmark, CRC-inhouse, and Her2ST), the self-attention strategy improved the Pearson correlation by 38.58%, 26.06%, and 20.39%, respectively, compared to the average performance of three patch-level single models. Similarly, for the TCGA-BRCA, TCGA-NSCLC, and TCGA-CRC WSI-level bulk gene expression prediction tasks, the Pearson correlation increased by 14.36%, 9.27%, and 42.55%, respectively, compared to the average performance of two WSI-level single models. By leveraging multiple pathological foundation models using Meta-Encoder, it can further improve molecular characterization in pathology images to advance precision oncology.

## Main text

Advances in artificial intelligence have significantly enhanced computational pathology, offering powerful tools for diagnosis and precision oncology[1]. Relevant tasks encompass tumor classification[2, 3], molecular biomarker prediction[4, 5], RNA expression prediction[6, 7], and more. The foundation models based on self-supervised learning strategies further revolutionize this field by enabling the effective utilization of large-scale unlabeled pathology images. In the past year, several patch-level pathological foundation models[8-13] and whole slide imaging (WSI)-level foundation models[8, 9, 14, 15] have been proposed. Despite their promising performance, different foundation models exhibit task-specific superiority, complicating appropriate model selection for targeted applications. Additionally, their divergent architectures and heterogeneous data sources complicate centralized retraining, particularly when data privacy constraints prohibit cross-institutional sharing of raw histopathology data.

However, publicly available model parameters open avenues for feature integration during fine-tuning. Inspired by meta-analysis, a statistical method that combines results from independent studies to improve the precision and efficiency of treatment effect estimation [16], we propose a Meta-Encoder framework. This framework integrates features extracted from multiple independent foundation models to produce a more comprehensive representation. This raises an important question: can features integrated through the Meta-Encoder, leveraging more diverse pretraining data, enhance downstream fine-tuning performance, and what is the optimal feature combination strategy for achieving peak performance? To explore this question, we performed a sequence of comparative experiments with increasing difficulty levels.

As shown in Figure 1a, we consider two major categories of foundation models: patch-level and WSI-level. For patch-level models, which extract features from individual patches, we selected three representative models published in 2024: CHIEF[8], GigaPath[9], and UNI[10]. CHIEF is pretrained on 60,530 WSIs spanning 19 anatomical sites from 14 cohorts. GigaPath is pretrained on 171,189 WSIs across 31 major tissue types from 28 cancer centers. UNI is pretrained on 100,426 WSIs covering 20 major tissue types from two hospitals and the Genotype-Tissue Expression (GTEx) consortium. These models were chosen because they were pretrained on datasets that do not overlap with the fine-tuning task datasets we tested, ensuring no data leakage and maintaining the fairness of the comparisons. While CHIEF and GigaPath are capable of producing WSI-level features in addition to their patch-level outputs, they show limited representational power for downstream tasks. Specialized WSI-level models are designed to extract comprehensive features from whole slide images. We selected two models specifically designed and trained for whole-slide feature representation: TITAN[14] and PRISM[15]. TITAN is a multimodal foundation model trained on 335,645 WSIs using visual self-supervised learning and vision-language alignment with pathology reports, along with 423,122 synthetic captions generated by a multimodal AI copilot. PRISM is pretrained on 587,000 WSIs with 195,000 clinical text reports.

**Figure 1:**
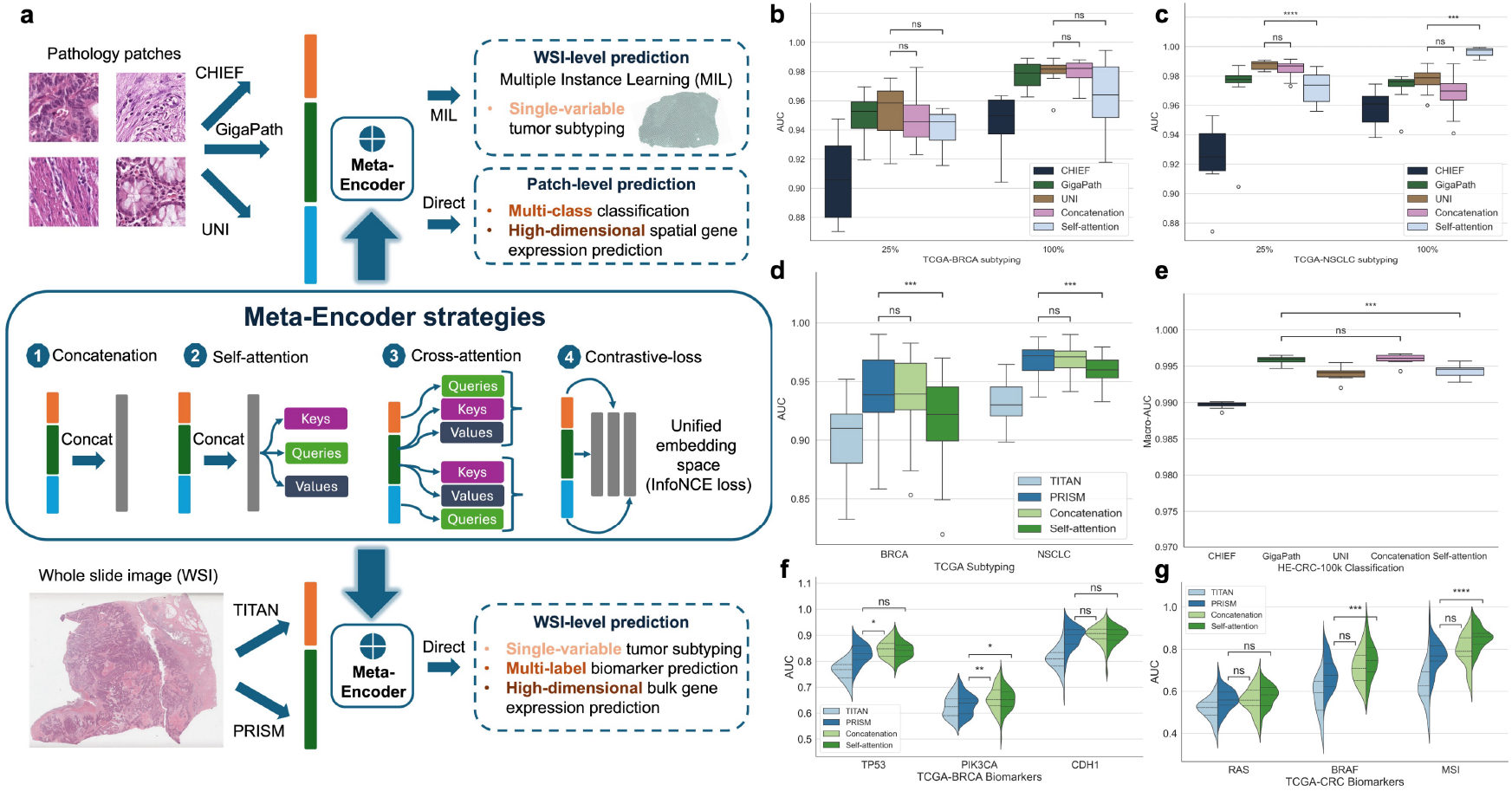
Illustration of Meta-Encoder strategies and their performance comparison across single-variable subtyping, multi-class classification, and multi-label tasks. (a) Overview of this study on integrating features via Meta-Encoder from various patch-level and WSI-level pathological foundation models using different strategies tailored for diverse downstream patch-level and WSI-level prediction tasks. (b-c) Box plots comparing single patch-level foundation models and various Meta-Encoder strategies for CLAM-based WSI-level subtyping of the TCGA-BRCA and NSCLC cohorts, where statistical significance was assessed using the t-test. The labels “25%” and “100%” indicate the percentage of the training dataset used for model training. (d) Box plots illustrating the performance of single WSI-level foundation models and different Meta-Encoder strategies in WSI-level subtyping of the TCGA-BRCA and NSCLC cohorts, with statistical testing performed using the t-test. (e) Box plots depicting the comparative performance of single patch-level foundation models and Meta-Encoder strategies in the 9-class patch classification task on the HE-CRC-100K dataset, with significance evaluated through the t-test. (f-g) Violin plots comparing the performance of single WSI-level foundation models and different Meta-Encoder strategies for WSI-level key biomarker prediction in the TCGA-BRCA and CRC cohorts, with statistical analysis conducted using the t-test.

Fine-tuning tasks can be stratified by increasing complexity based on the dimensionality of the target variable *Y*, ranging from binary tumor subtyping, to multi-class classification, multi-label biomarker prediction, and ultimately high-dimensional gene expression prediction. Remarkably, these tasks span both the patch-level and WSI-level. For WSI-level foundation models, the extracted features can be directly used for fine-tuning on downstream WSI-level tasks. On the other hand, for patch-level foundation models, the extracted features can be directly applied to downstream patch-level tasks. However, for WSI-level predictions using patch-level models, multiple instance learning (MIL) framework such as clustering-constrained-attention multiple-instance learning (CLAM)[17] should be employed to aggregate patch-level features effectively.

We adopted four Meta-Encoder strategies for feature fusion across multiple pathological foundation models. Two of these are relatively basic: the concatenation strategy and the self-attention strategy. In the concatenation strategy, features extracted from different foundation models for each patch are concatenated and then used for subsequent training. For the self-attention strategy, features are first concatenated and then passed through a self-attention layer, while the remaining pipeline is consistent with the single model and concatenation-based fusion approaches. Building on these, we employed two more advanced strategies. One approach is the cross-attention strategy, where features extracted from different models are treated as Key, Query, and Value, enabling the cross-attention mechanism to capture high-order interactions between them. Another approach is the contrastive-loss strategy, which is based on the idea of treating features extracted by different models from the same image as complementary descriptions of the same object from different perspectives. These features are linearly mapped into a unified embedding space, where positive (similar features) and negative (dissimilar features) pairs are formed to compute the contrastive loss. The objective is to minimize the distance between features from the same image in this common space while maximizing the distance between features from different images.

For single-variable subtype classification tasks, the results achieved with Meta-Encoder feature fusion can rival those of the best single model, as detailed in the following experiments. In the context of CLAM-based patch-level tumor subtyping, where the prediction tasks involve binary classification for The Cancer Genome Atlas Program (TCGA) breast cancer (BRCA) and non-small cell lung cancer (NSCLC), the UNI foundation model consistently achieves the best performance. This may be attributed to its shared origin with the CLAM framework, both developed by the same team, which enhances their architectural compatibility. Feature concatenation of all three foundation models followed by downstream fine-tuning performs comparably to UNI, without notable improvement (Figure 1b, c, and Extended Data Table S1). The self-attention fusion strategy performs comparably to UNI in BRCA data, worse with 25% NSCLC data, and better with 100% NSCLC data, but overall fails to achieve consistent gains (Figure 1b, c, and Extended Data Table S1). With respect to WSI-level foundation models, both feature concatenation and the best-performing individual foundation model achieve strong subtyping performance, while the introduction of non-linear interactions via self-attention slightly degrades performance (Figure 1d and Extended Data Table S2). Overall, in single-variable prediction settings, feature concatenation performs on par with the best individual foundation model. Despite the strong performance of individual models on single-variable binary classification tasks, the feature fusion strategy such as concatenation via the Meta-Encoder, matching the best individual models, still retains practical value in these simple tasks, which can ensure robust performance and reduce the time required for model selection and tuning. For the 9-class classification task on the HE-CRC-100K dataset, features from individual patch-level foundation models already exhibit strong discriminative power, with concatenation and self-attention strategies maintaining comparable performance (Figure 1e and Extended Data Table S3). Notably, UNI achieves the best performance among single models on the subtyping task, while GigaPath performs best on this multi-class classification task, indicating task-dependent strengths of individual models. Nevertheless, feature fusion via the Meta-Encoder can enhance the performance lower bound across models. For the multi-label classification, feature concatenation improves the average performance across multiple biomarker predictions (Extended Data Table S4). While implementing self-attention to capture inter-model feature relationships yields no additional gain over concatenation in TCGA-BRCA (Figure 1f), it leads to a notable improvement in TCGA-CRC (Figure 1g). This may relate to task difficulty, as these three biomarkers in CRC are more challenging to predict than those in BRCA, making the modeling of inter-model feature interactions via self-attention more beneficial. In summary, for tasks of moderate complexity, the concatenation strategy via Meta-Encoders is a reliable approach. Particularly for new tasks, using a Meta-Encoder can ensure a robust performance baseline while saving the time and effort required to identify the optimal single model.

For complex tasks with high-dimensional prediction targets, the Meta-Encoder demonstrates more pronounced advantages. The HEST-Benchmark comprises nine tasks aimed at predicting the spatial expression of the top 50 genes with the highest variance of different cancer types[18], representing higher-dimensional prediction targets compared to previous downstream fine-tuning tasks. Feature concatenation yields noticeable improvements over individual foundation models in spatial gene expression prediction. Further gains are achieved by introducing non-linearity through the self-attention strategy and imposing constraints in the latent space via the contrastive-loss strategy, while the cross-attention strategy performs comparably to simple concatenation (Figure 2a and Extended Data Table S5). For the other two spatial transcriptomics datasets, CRC-inhouse and Her2ST, feature concatenation provides improvements over single foundation models for predicting spatial gene expression (Figure 2b, c, and Extended Data Table S6). Self-attention, which introduces non-linearity, performs comparably to concatenation. Although the cross-attention strategy outperforms single models, its performance drops notably relative to the concatenation strategy, potentially due to its complexity hindering effective fine-tuning. Contrastive-loss strategy via Meta-Encoder achieves the best accuracy but requires substantially longer training time, with only marginal performance gains, raising concerns about its efficiency and practical utility (Extended Data Table S7). Regarding bulk gene expression prediction based on WSI-level features, we utilized WSIs from TCGA to predict corresponding bulk RNA-seq profiles, focusing on highly variable genes identified from spatial transcriptomics data for each cancer type. Feature concatenation via Meta-Encoder shows clear gains for BRCA, NSCLC, and CRC (Figure 2d, e, f). In BRCA and CRC, Meta-Encoder strategies including self-attention, cross-attention and contrastive-loss can further improve performance, whereas in NSCLC, these Meta-Encoder strategies perform similarly to concatenation (Extended Data Table S8). Experimental results demonstrate that Meta-Encoder-based feature fusion consistently enhances performance compared to single foundation models in high-dimensional prediction tasks. For such tasks, we particularly recommend using the self-attention strategy via Meta-Encoder. For three patch-level spatial gene expression prediction tasks (HEST-Benchmark, CRC-inhouse, and Her2ST), this strategy improved Pearson correlation by 0.079, 0.055, and 0.032, corresponding to relative gains of 38.58%, 26.06%, and 20.39%, respectively, over the average performance of three patch-level single models. Similarly, on the WSI-level bulk gene expression prediction tasks (TCGA-BRCA, TCGA-NSCLC, and TCGA-CRC), the self-attention strategy increased Pearson correlation by 0.052, 0.036, and 0.060, with relative improvements of 14.36%, 9.27%, and 42.55%, respectively, compared to the average performance of two WSI-level single models.

**Figure 2:**
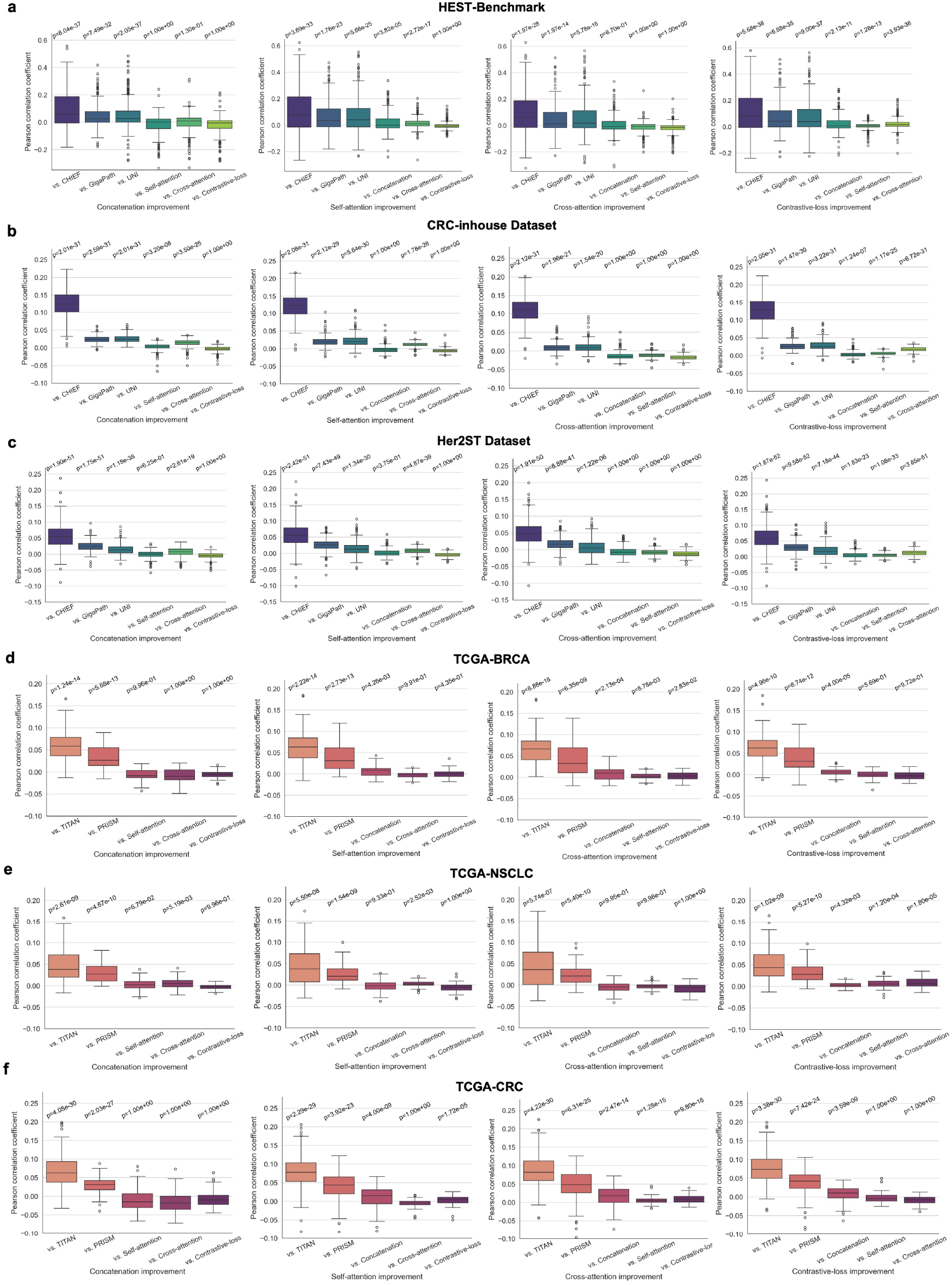
Comparative performance analysis of single foundation models and different Meta-Encoder strategies for high-dimensional gene expression prediction tasks. (a-c) Box plots illustrating gene-specific improvements in Pearson correlation across various single patch-level foundation models and Meta-Encoder strategies for high-dimensional spatial gene expression prediction in HEST-Benchmark, CRC-inhouse, and Her2ST datasets, where each point in the box plot represents the Pearson correlation of each gene predicted by the Meta-Encoder strategy on the x-axis minus the Pearson correlation of the corresponding gene from the vs. model, with statistical significance assessed using the one-sided Wilcoxon signed-rank test. (d-f) Box plots depicting gene-specific improvements in Pearson correlation across various single WSI-level foundation models and Meta-Encoder strategies for high-dimensional bulk gene expression prediction across TCGA-BRCA, NSCLC, and CRC cohorts.

Several studies have attempted to integrate multiple pathological foundation models. For instance, Ma et al. proposed a Generalizable Pathology Foundation Model (GPFM) by employing unified knowledge distillation to combine multiple foundation models in the field of pathology[19]. However, this approach requires repeating large-scale pretraining process, utilizing 190 million images from approximately 86,000 public HE-stained WSIs covering 34 major tissue types. Additionally, Neidlinger et al. introduced Efficient Approach for Guided Local Examination (EAGLE)[20], which integrates two foundation models: CHIEF, for efficient tile selection, and Virchow2[21], for extracting high-quality patch-level features. This method essentially combines the weights of patches from WSI-level foundation models with patch features from patch-level foundation models. In contrast, the focus of our Meta-Encoder framework is on integrating features from different foundation models in downstream fine-tuning tasks. This approach is more lightweight and generalizable, thereby offering greater flexibility for various applications.

Our systematic experiments reveal that feature fusion via Meta-Encoder effectively leverages the complementary strengths of multiple foundation models pretrained on diverse datasets, with increasing benefits in more complex tasks. For simple single-variable tasks, fused features perform similarly to or slightly worse than the best single model. However, we still recommend using the concatenation strategy via Meta-Encoder to reduce the time and effort required for model selection and tuning. For moderately complex tasks like multi-class or multi-label classification, Meta-Encoder-based concatenation and self-attention strategies improve performance. Concatenation is sufficient for low-complexity tasks, while self-attention is better for more challenging ones. For highly complex tasks, such as high-dimensional gene prediction, feature fusion significantly improves performance. We tested advanced fusion strategies like cross-attention and contrastive-loss in addition to concatenation and self-attention. While cross-attention and contrastive-loss occasionally outperform self-attention, their overall improvements are limited. Moreover, cross-attention introduces structural complexity, especially when fusing three or more models, as multiple KQV combinations and selection increase subjectivity. The contrastive-loss strategy also incurs substantial computational overhead due to the requirement of calculating contrastive loss during each batch optimization. Therefore, we recommend self-attention as the most practical and effective Meta-Encoder strategy for high-dimensional gene expression prediction, which can further facilitate the molecular characterization of large cancer cohorts that only have pathological images but lack omics data, thereby advancing prognosis prediction[22].

In summary, do we need feature fusion across multiple pathological foundation models via Meta-Encoder for downstream fine-tuning tasks? The answer is: yes! Overall, our Meta-Encoder framework proves highly effective for downstream fine-tuning tasks. Which feature fusion strategy yields optimal performance? For low-complexity tasks, the concatenation strategy can achieve performance comparable to the best single model, thus saving time on model selection and tuning. In moderately complex multi-class or multi-label tasks, the concatenation strategy maintained strong performance, while the self-attention strategy showed more benefits in highly challenging scenarios.

Looking ahead, the development of foundation models in the medical field should focus less on repetitive efforts for similar tasks and more on effective resource integration to build a more comprehensive and synergistic ecosystem. Furthermore, the increasing significance of multimodal approaches represents a critical emerging trend. HE-stained pathology images capture rich histological and morphological features, spatial transcriptomics provide spatial RNA-level characteristics, and multiplex immunohistochemistry (mIHC) highlights spatial protein expression. These modalities are complementary and collectively provide a more complete representation of physiological and pathological states. Our Meta-Encoder framework also holds promise for the tight integration of multimodal pathology foundation models, combining morphological characteristics with molecular-level features to further advance the clinical adoption of computational pathology.

## Methods

### TCGA tumor subtyping

For tumor subtyping prediction on WSIs from the TCGA dataset, two approaches can be applied: (1) using patch-level foundation models to extract features, followed by aggregating features from all patches within a WSI via multiple instance learning (MIL) for prediction; or (2) directly using WSI-level foundation models to extract WSI features for downstream fine-tuning. The TCGA-BRCA and TCGA-NSCLC datasets were used in this study. Classification predictions were conducted on TCGA-BRCA to differentiate between invasive ductal carcinoma (IDC) and invasive lobular carcinoma (ILC) tumor subtypes in 837 breast cancer patients, and on TCGA-NSCLC to distinguish between lung adenocarcinoma (LUAD) and lung squamous cell carcinoma (LUSC) in 862 non-small cell lung cancer (NSCLC) patients (Extended Data Table S9). For the patch-level features fusion for tumor subtyping, we utilized three patch-level foundation models to extract patch features, followed by downstream fine-tuning using the CLAM framework. For foundation models capable of processing entire WSIs, WSI-level feature representations were directly obtained for downstream fine-tuning.

### Multi-class classification for patches on the HE-CRC-100K dataset

The HE-CRC-100K dataset contains patches from nine distinct labels (Extended Data Table S10). These patches were manually extracted from 86 HE-stained human cancer tissue slides derived from formalin-fixed paraffin-embedded (FFPE) samples. Patch-level features are extracted using foundation models and then mapped to a 9-dimensional vector via a linear layer for multi-class classification.

The dataset has a predefined train/test split. To ensure robustness, each experiment is repeated ten times, and the mean and standard deviation of performance metrics are reported. The learning rate is scheduled using cosine annealing, with an initial value of 1e-4 and a minimum of 1e-6 over five cycles. The batch size is set to 512, and an infinite data loader is employed, with a total of 10,000 training iterations. Cross-entropy loss is used as the loss function. The training and evaluation processes are repeated ten times to compute the mean and standard deviation for performance assessment.

### Multi-Label prediction of tumor biomarkers using WSI-level representations

Directly predicting the mutation status of key genes and molecular biomarkers from WSIs holds significant clinical value. As multiple genetic alterations and biomarkers are of interest, this task is naturally formulated as a multi-label prediction problem. For the TCGA-BRCA cohort, we focus on simultaneously predicting the mutation status of three frequently mutated genes: *TP53, PIK3CA*, and *CDH1*. For the TCGA-CRC cohort, we target clinically relevant mutations and biomarkers closely associated with treatment decisions, including *RAS* (*KRAS*/*NRAS*) mutations, *BRAF* mutations, and microsatellite instability (MSI) status. The prevalence of these biomarkers is summarized in the Extended Data Table S11.

### High-dimensional spatial gene expression prediction using patch-level features

The HEST-Benchmark defines nine tasks using data from eight human cancer types across nine organs, including eight primary and one metastatic datasets. These tasks are designed to evaluate the ability of different models to predict spatial gene expression from histology image patches (Extended Data Table S12). For each task, we predict the expression levels of the top 50 genes with the highest normalized variance across all samples, using HE image regions measuring 112×112 μ*m*, which correspond to 224×224 pixels at 20× magnification. To avoid data leakage at the patient level, we adopt patient-stratified *k*-fold cross-validation, where *k* equals the number of patients. For the clear cell renal cell carcinoma (ccRCC) task, *k* is halved to accommodate the large number of patients. For the expression of target genes in each spot, we first preprocess the data by normalizing the gene counts by the total gene counts, followed by applying *log*(1 + *p*), which computes the natural logarithm of one plus the input value.

We further constructed patch-spot pairs on two additional spatial transcriptomics datasets to evaluate the performance of different methods in predicting gene spatial expression. The CRC-inhouse dataset comprises ten 10x Visium spatial transcriptomics slides from ten CRC patients, while the Her2ST dataset consists of 36 legacy ST slides from eight Her2+ breast cancer patients (Extended Data Table S13). Following the same pipeline as in the HEST-Benchmark construction, we applied a robust tissue-versus-background detection method based on a DeepLabV3 model[23] with an ImageNet-pretrained ResNet50 backbone, fine-tuned on a set of manually annotated segmentation masks. After segmentation, 42,123 patch-spot pairs were generated for the CRC-inhouse dataset and 13,595 for the Her2ST dataset.

### High-dimensional bulk gene expression prediction based on WSI-level features

The samples from TCGA with both WSIs and bulk RNA-seq data were used for training and testing downstream bulk gene expression prediction tasks, including the BRCA, NSCLC, and CRC cohorts. As detailed in Extended Data Table S14, the prediction target in the BRCA cohort is 50 genes with highest variance from the HEST breast cancer ST dataset. For the NSCLC cohort, the prediction targets 50 genes with highest variance from the HEST lung cancer ST dataset. In the CRC cohort, the prediction focuses on genes with high variance from the in-house CRC ST dataset, with 174 genes detected in the bulk RNA-seq of the TCGA-CRC cohort.

## Data availability

All clinical features, WSI images and RNA-seq data from the public dataset TCGA can be downloaded from https://portal.gdc.cancer.gov. The preprocessed molecular features for TCGA used in our experiments are available at https://github.com/KatherLab/cancer-metadata. The public dataset HE-CRC-100K used for multi-class classification can be downloaded from https://zenodo.org/record/1214456. The HEST-Benchmark used for patch-level high-dimensional spatial gene expression prediction can be downloaded from https://huggingface.co/datasets/MahmoodLab/hest-bench. The CRC-inhouse dataset used in this study is available from the original publication https://www.cell.com/cell-reports-medicine/fulltext/S2666-3791(24)00205-2. The Her2ST dataset can be downloaded from https://zenodo.org/records/3957257.

## Code availability

All implementation codes related to this study have been made openly accessible via our GitHub repository: https://github.com/ruitian-olivia/Meta-Encoder.

