## Supplementary Information for "Features fusion or not: harnessing multiple pathological foundation models using Meta-Encoder for downstream tasks fine-tuning"

Table S1. Performance comparison of single patch-level foundation models and feature fusion via Meta-Encoder in CLAM-based patch-level tumor subtyping

| Model type | BRCA subtype AUC (25%) | BRCA subtype AUC (100%) | NSCLC subtype AUC (25%) | NSCLC subtype AUC (100%) |
|---|---|---|---|---|
| CHIEF | 0.9070±0.0282 | 0.9454±0.0190 | 0.9250±0.0225 | 0.9582±0.0121 |
| GigaPath | <u>0.9502±0.0153</u> | 0.9773±0.0093 | 0.9714±0.0238 | 0.9722±0.0111 |
| UNI | **0.9519±0.0212** | **0.9795±0.0101** | **0.9873±0.0031** | <u>0.9772±0.0088</u> |
| Concatenation | 0.9489±0.0194 | <u>0.9794±0.0087</u> | <u>0.9847±0.0063</u> | 0.9672±0.0137 |
| Self-attention | 0.9398±0.0147 | 0.9638±0.0243 | 0.9724±0.0109 | **0.9964±0.0030** |

Table S2. Performance comparison of single WSI-level foundation models and feature fusion via Meta-Encoder in WSI-level tumor subtyping

| Model type | BRCA subtype AUC | NSCLC subtype AUC |
|---|---|---|
| TITAN | 0.9019±0.0288 | 0.9312±0.0168 |
| PRISM | **0.9419±0.0290** | <u>0.9678±0.0117</u> |
| Concatenation | <u>0.9418±0.0287</u> | **0.9688±0.0109** |
| Self-attention | 0.9205±0.0319 | 0.9592±0.0109 |

Table S3. Performance comparison of single patch-level foundation models and feature fusion via Meta-Encoder in 9-class patch classification on HE-CRC-100K

| Model type | HE-CRC-100k 9-type classification macro-AUC |
|---|---|
| CHIEF | 0.9896±0.0004 |
| GigaPath | <u>0.9958±0.0006</u> |
| UNI | 0.9939±0.0009 |
| Concatenation | **0.9960±0.0007** |
| Self-attention | 0.9944±0.0009 |

Table S4. Performance comparison of single WSI-level foundation models and feature fusion via Meta-Encoder in multi-label biomarker prediction for TCGA cohorts

| Cohort | TCGA-BRCA | | | | TCGA-CRC | | | |
|---|---|---|---|---|---|---|---|---|
| Biomarker | TP53 mutation | PIK3CA mutation | CDH1 mutation | Mean AUC | RAS mutation | BRAF mutation | MSI status | Mean AUC |
| TITAN | 0.7623±0.0354 | 0.6242±0.0425 | 0.8084±0.0456 | 0.7316 | 0.5141±0.0517 | 0.5801±0.0925 | 0.6340±0.0942 | 0.5761 |
| PRISM | 0.8284±0.0390 | 0.6329±0.0397 | 0.8936±0.0411 | 0.7850 | <u>0.5611±0.0513</u> | 0.6837±0.0742 | 0.7677±0.0695 | 0.6708 |
| Concatenation | **0.8467±0.0313** | **0.6558±0.0442** | **0.9020±0.0369** | **0.8015** | 0.5588±0.0619 | <u>0.7133±0.0748</u> | <u>0.7934±0.0756</u> | <u>0.6885</u> |
| Self-attention | <u>0.8415±0.0316</u> | <u>0.6508±0.0454</u> | <u>0.8981±0.0355</u> | <u>0.7968</u> | **0.5757±0.0516** | **0.7464±0.0836** | **0.8546±0.0507** | **0.7256** |

Table S5. Performance comparison of single patch-level foundation models and feature fusion via Meta-Encoder in spatial gene prediction across nine HEST-Benchmark tasks

| Model type | HEST-Benchmark Pearson Correlation | | | | | | | | | |
|---|---|---|---|---|---|---|---|---|---|---|
| | CCRCC | COAD | IDC | LUAD | LYMPH-IDC | PAAD | PRAD | READ | SKCM | Mean |
| CHIEF | 0.1219±0.0745 | 0.1196±0.0555 | 0.2608±0.0700 | 0.2464±0.0548 | **0.2519±0.0540** | 0.1249±0.1064 | <u>0.2252±0.1159</u> | 0.0649±0.0477 | 0.1516±0.0583 | 0.1741 |
| GigaPath | 0.1342±0.0782 | 0.1751±0.0544 | 0.3331±0.1029 | 0.3146±0.0283 | 0.2250±0.0670 | 0.2297±0.0641 | **0.2260±0.0902** | 0.1757±0.0313 | 0.2133±0.0546 | 0.2252 |
| UNI | <u>0.1415±0.0785</u> | 0.1633±0.0411 | 0.3569±0.0970 | 0.3108±0.0229 | 0.2228±0.0570 | 0.1986±0.0850 | 0.2033±0.0904 | 0.1540±0.0324 | 0.1866±0.0379 | 0.2153 |
| Concatenation | **0.1419±0.0781** | 0.1991±0.0556 | 0.3831±0.0946 | 0.4062±0.0392 | <u>0.2274±0.0555</u> | 0.3123±0.0375 | 0.2161±0.0915 | 0.1954±0.0219 | 0.3236±0.0392 | 0.2672 |
| Self-attention | 0.1267±0.0687 | <u>0.2373±0.0428</u> | <u>0.3876±0.0996</u> | **0.4647±0.0343** | 0.2071±0.0427 | <u>0.3654±0.0604</u> | 0.1740±0.0810 | <u>0.1962±0.0341</u> | **0.3966±0.0309** | <u>0.2839</u> |
| Cross-attention | 0.1228±0.0764 | 0.2203±0.0597 | 0.3543±0.1092 | 0.4323±0.0108 | 0.2064±0.0467 | 0.3534±0.0441 | 0.1867±0.0859 | 0.1906±0.0380 | 0.3557±0.0167 | 0.2692 |
| Contrastive-loss | 0.1351±0.0727 | **0.2442±0.0434** | **0.3971±0.0988** | <u>0.4549±0.0497</u> | 0.2220±0.0440 | **0.3689±0.0531** | 0.1881±0.0894 | **0.2052±0.0272** | <u>0.3850±0.0445</u> | **0.2889** |

Table 6. Performance comparison of single patch-level foundation models and feature fusion via Meta-Encoder in spatial gene prediction for CRC-inhouse and Her2ST dataset

| Model type | Pearson Correlation | |
|---|---|---|
| | CRC-inhouse | Her2ST |
| CHIEF | 0.1445±0.0670 | 0.1338±0.0760 |
| GigaPath | 0.2438±0.0837 | 0.1649±0.0931 |
| UNI | 0.2421±0.0773 | 0.1755±0.1096 |
| Concatenation | <u>0.2676±0.0805</u> | 0.1894±0.1163 |
| Self-attention | 0.2649±0.0711 | <u>0.1903±0.1277</u> |
| Cross-attention | 0.2537±0.0739 | 0.1826±0.1187 |
| Contrastive-loss | **0.2710±0.0753** | **0.1952±0.1278** |

Table S7. Training time comparison of single patch-level foundation models and feature fusion via Meta-Encoder in spatial gene prediction tasks

| Model type | Training time (s) / epochs | | |
|---|---|---|---|
| | HEST-Benchmark | CRC-inhouse | Her2ST |
| CHIEF | 1.602 | 3.030 | 1.850 |
| GigaPath | 1.528 | 2.995 | 1.925 |
| UNI | 1.390 | 2.980 | 1.638 |
| Concatenation | 1.331 | 2.915 | 2.275 |
| Self-attention | 1.635 | 3.390 | 1.975 |
| Cross-attention | 1.582 | 3.375 | 2.300 |
| Contrastive-loss | 24.394 | 38.850 | 13.925 |

Table S8. Performance comparison of single WSI-level foundation models and feature fusion via Meta-Encoder in bulk gene expression prediction for TCGA cohorts

| Model type | Pearson correlation | | |
|---|---|---|---|
| | TCGA-BRCA | TCGA-NSCLC | TCGA-CRC |
| TITAN | 0.3455±0.0145 | 0.3761±0.0212 | 0.1222±0.0277 |
| PRISM | 0.3733±0.0196 | 0.3945±0.0217 | 0.1605±0.0229 |
| Concatenation | 0.4048±0.0160 | <u>0.4241±0.0200</u> | 0.1902±0.0260 |
| Self-attention | <u>0.4110±0.0162</u> | 0.4210±0.0221 | <u>0.2015±0.0250</u> |
| Cross-attention | **0.4136±0.0156** | 0.4184±0.0212 | **0.2068±0.0251** |
| Contrastive loss | 0.4107±0.0166 | **0.4269±0.0206** | 0.1982±0.0254 |

Table S9. Description of TCGA datasets utilized in tumor subtyping

| Cohorts | Subtypes | Number (percentage) |
|---|---|---|
| TCGA-BRCA (N=837) | IDC | 694(82.92%) |
| | ILC | 143 (17.08%) |
| TCGA-NSCLC (N=862) | LUAD | 432(50.12%) |
| | LUSC | 430(49.88%) |

Table S10. Category counts and proportions in the HE-CRC-100K dataset

| HE-CRC-100K label | Training set | Test set |
|---|---|---|
| Adipose (ADI) | 10,407(10.41%) | 1,338(18.64%) |
| Background (BACK) | 10,566(10.57%) | 847(11.80%) |
| Debris (DEB) | 11,512(11.51%) | 339(4.72%) |
| Lymphocytes (LYM) | 11,557(11.56%) | 634(8.83%) |
| Mucus (MUC) | 8,896(8.9%) | 1,035(14.42%) |
| Smooth muscle (MUS) | 13,536(13.54%) | 592(8.25%) |
| Normal colon mucosa (NORM) | 8,763(8.76%) | 741(10.32%) |
| Cancer-associated stroma (STR) | 10,446(10.45%) | 421(5.86%) |
| Colorectal adenocarcinoma epithelium (TUM) | 14,317(14.32%) | 1,233(17.17%) |

Table S11. Prevalence of target biomarkers in TCGA-BRCA and TCGA-CRC

| Cohort | TCGA-BRCA (N=887) | | | TCGA-CRC (N=442) | | |
|---|---|---|---|---|---|---|
| Biomarker | *TP53* | *PIK3CA* | *CDH1* | *RAS* (*KRAS*+*NRAS*) | *BRAF* | MSI |
| Mutation /Positive | 314 (35.40%) | 311 (35.06%) | 108 (12.18%) | 224 (50.68%) | 57 (12.90%) | 64 (14.48%) |
| Wildtype /Negative | 573 (64.60%) | 576 (64.94%) | 779 (87.82%) | 218 (49.32%) | 385 (87.10%) | 378 (85.52%) |

Table S12. Data description of nine tasks in the HEST-Benchmark

| Task name | Cancer type | Patch-spot number | Tissue number | Patient number | Fold number | Target genes number |
|---|---|---|---|---|---|---|
| Clear cell renal cell carcinoma (ccRCC) | Kidney cancer | 74,220 | 24 | 12 | 6 | 50 |
| colonic adenocarcinoma (COAD) | Colon cancer | 15,651 | 4 | 2 | 2 | 50 |
| invasive ductal carcinoma (IDC) | Breast cancer | 35,536 | 4 | 4 | 4 | 50 |
| lung adenocarcinoma (LUAD) | Lung cancer | 5,206 | 2 | 2 | 2 | 50 |
| axillary lymph nodes in IDC (LYMPH-IDC) | Metastatic of breast cancer | 19,964 | 4 | 4 | 4 | 50 |
| pancreatic adenocarcinoma (PAAD) | Pancreatic cancer | 7,571 | 3 | 3 | 3 | 50 |
| prostate adenocarcinoma (PRAD) | Prostate cancer | 62,710 | 23 | 2 | 2 | 50 |
| rectal adenocarcinoma (READ) | Rectum cancer | 8,407 | 4 | 2 | 2 | 50 |
| skin cutaneous melanoma (SKCM) | Skin cancer | 3,034 | 2 | 2 | 2 | 50 |

Table S13. Description of the CRC-Inhouse and Her2ST Datasets

| Dataset | Cancer type | Patch-spot number | Tissue number | Patient number | Fold number | Target gene number |
|---|---|---|---|---|---|---|
| CRC-inhouse (colorectal cancer) | Colon cancer & Rectum cancer | 42,123 | 10 | 10 | 10 | 179 |
| Her2ST (Her2+ breast cancer) | Breast cancer | 13,595 | 36 | 8 | 8 | 321 |

Table S14. Description of TCGA cohorts for bulk gene expression prediction

| Cohort | TCGA-BRCA (N=1,059) | TCGA-NSCLC (N=949) | TCGA-CRC (N=609) |
|---|---|---|---|
| Target gene number | 50 | 50 | 174 |

### Supplementary Methods

#### Fine-tuning details of TCGA tumor subtyping

For the CLAM-based patch-level features fusion for tumor subtyping, we utilized three patch-level foundation models to extract patch features, followed by downstream fine-tuning using the CLAM framework. This framework leverages attention mechanisms to automatically identify diagnostically significant subregions, enabling accurate classification of entire whole slide images (WSIs). In this setup, we randomly split the data ten times, with 10% as the validation set, 10% as the test set, and the remaining 80% as the training set. To simulate a more data-limited scenario, the dataset was downsampled to 25% of its original size, followed by an 8:1:1 split. For downstream training with individual foundation model features, we input the extracted CHIEF, GigaPath, and UNI feature vectors into the CLAM framework. The training process was configured with a maximum of 20 epochs, a patience of 10, an instance-level clustering loss function based on SVM, a slide-level classification loss function using cross-entropy, and a learning rate of 2e-4. The encoding sizes for UNI, GigaPath, and CHIEF are 1024, 1536, and 768, respectively. Each is mapped to a 512-dimensional space via a linear layer, followed by a ReLU-activated multilayer perceptron (MLP). An attention network with sigmoid gating, consisting of three sequential fully connected layers with dimensions 512, 256, and 1, is applied to generate an attention score of shape $N \times 1$, which is subsequently used to weight the feature matrix. A softmax operation is subsequently applied to yield the final output. The model is trained for a maximum of 20 epochs with an early stopping patience of 10. During the MIL process, the batch size is set to 1. In the concatenation strategy, features extracted from three foundation models for each patch are concatenated and input into the CLAM framework, resulting in a combined encoding size of 3328. The downstream architecture mirrors that of the single-model CLAM setup. For the self-attention strategy, features are first concatenated and passed through a self-attention layer (embedding dimension = 1024, number of heads = 1). The subsequent pipeline remains consistent with that of the single-model and concatenation-based fusion approaches.

In WSI-level feature fusion for tumor subtyping, WSI-level feature representations are directly extracted from foundation models capable of processing whole slide images, followed by downstream fine-tuning. Model performance is evaluated via five-fold cross-validation, repeated ten times with different random seeds. WSI features are extracted using TITAN and PRISM, yielding feature dimensions of 768 and 1280, respectively. To assess the linear separability of these representations, a single-layer linear classifier followed by a sigmoid activation is applied. In the concatenation strategy, features extracted from each WSI by different foundation models are concatenated into a 2048-dimensional vector, which is then used to fine-tune a downstream classification model. For the self-attention strategy, the concatenated features are first passed through a self-attention layer (embedding dimension = 1024, number of heads = 8), followed by a linear classification layer. In five-fold cross-validation, four folds are used for training and the remaining fold for testing, without a separate validation set. To prevent overfitting and facilitate better convergence in the later stages of training, the learning rate is dynamically adjusted using cosine annealing schedule. The learning rate decays from an initial value of 5e-4 to a minimum of 1e-5 following a cosine schedule, completed within a single annealing cycle (cycle count = 1). The model is optimized using the Adam optimizer, with a total of 100 training epochs and a batch size of 128. Focal loss is employed as the loss function, which extends binary cross-entropy by incorporating a modulating factor to focus training on hard examples, making it suitable for handling class imbalance. For the breast cancer binary classification task, where the class distribution between IDC and ILC is imbalanced, the alpha parameter is set to 0.8 to reweight positive and negative samples. In the NSCLC subtype classification task, where class proportions are approximately balanced, alpha is set to 0.5. In both tasks, the focusing parameter gamma is set to 2.

**Fine-tuning details of multi-class classification for patches in the HE-CRC-100K dataset**

The HE-CRC-100K dataset has a predefined train/test split. To ensure robustness, each experiment is repeated ten times, and the mean and standard deviation of performance metrics are reported. The learning rate is scheduled using cosine annealing schedule, with an initial value of 1e-4 and a minimum of 1e-6 over five cycles. The batch size is set to 512, and an infinite data loader is employed, with a total of 10,000 training iterations. Cross-entropy loss is used as the loss function.

**Fine-tuning details of multi-Label prediction of tumor biomarkers using WSI-level representations**

All WSI data with mutation labels are partitioned using five-fold cross-validation, repeated ten times to generate ten distinct data splits. The learning rate follows a cosine annealing schedule, decaying from an initial value of 1e-4 to a minimum of 1e-6 within a single annealing cycle. The model is optimized using the Adam optimizer with 100 training epochs and a batch size of 256. Focal loss is employed as the objective function, with the focusing parameter gamma set to 2 across all tasks. For the BRCA cohort, the alpha parameters for predicting *TP53*, *PIK3CA*, and *CDH1* mutations are set to 0.6, 0.6, and 0.9, respectively. For the CRC cohort, alpha values for *RAS* and *BRAF* mutations and MSI status are set to 0.5, 0.8, and 0.8, respectively. Consistent with the WSI-level tumor subtyping setup, WSI features are extracted using TITAN and PRISM models to predict the status of multiple biomarkers. In the concatenation strategy, features from both models are combined before prediction. In the self-attention strategy, the concatenated features are first passed through a cross-attention layer (embedding dimension = 1024; number of heads = 8), followed by a linear layer for multi-label classification.

**Fine-tuning details of high-dimensional spatial gene expression prediction using patch-level features**

During the fine-tuning process on the HEST-Benchmark, the batch size is set to 512, with a total of 50 epochs. The learning rate follows cosine annealing schedule, starting from 1e-4 and decaying to 1e-6, with a single annealing cycle. The loss function used during training is the mean squared error (MSE). In the concatenation strategy, features are merged together. In the self-attention approach, the concatenated features are first mapped to a 1024-dimensional space, with 8 attention heads. For the cross-attention strategy, GigaPath features serve as the Key and Value, while UNI features are used as the Query, and the output is mapped to a 1024-dimensional space with 8 heads. CHIEF features are also used as the Query and mapped to 1024 dimensions. These two are then combined into a 2048-dimensional vector for target gene expression prediction. The concept behind the contrastive-loss strategy is to treat features extracted by different foundation models from the same image as descriptions of one object from different perspectives. These features are linearly mapped into a common space of 1024 dimensions, from which positive and negative pairs are formed to compute the contrastive loss. The goal is to bring features of the same image closer in this common space, while maximizing the distance between features from different images. Because contrastive loss is computed intra-batch, we set the batch size to 256 to meet GPU memory constraints. The total number of epochs is set to 25, ensuring the overall number of iterations remains comparable to other methods. The weight for contrastive loss is set to 0.1. More specifically, for the contrastive-loss strategy in Meta-Encoder framework, we adopt the information noise contrastive estimation (InfoNCE) loss[1], which encourages high similarity between positive pairs and low similarity between negative pairs in the embedding space. InfoNCE formulates contrastive learning as a classification task, where the model distinguishes a positive pair from a set of negative pairs. Similarity is computed using a softmax-based probabilistic framework.

During training on the CRC-inhouse and Her2ST datasets, the batch size was set to 512 and the number of epochs to 20. The learning rate followed a cosine annealing schedule, decreasing from 1e-4 to 1e-6 with an annealing cycle of 1. The Meta-Encoder strategies, including concatenation, self-attention, cross-attention, and contrastive-loss, were applied in a manner consistent with the approach used in the HEST-Benchmark. Regarding the contrastive-loss strategy, the unified embedding space was set to 1024 dimensions, with a batch size of 256 and 10 epochs. For the CRC-inhouse dataset, which involves predicting 179 genes, the contrastive loss was added to the MSE loss with a weight of 0.1. For the Her2ST dataset, which involves predicting 321 genes, the contrastive loss was weighted at 0.5.

**Fine-tuning details of high-dimensional bulk gene expression prediction based on WSI-level features**

The STAR-processed mRNA count data were downloaded from TCGA, and the FPKM-UQ data were selected for use. The data were preprocessed using $log\,2(FPKM\_UQ + 1)$ transformation to approximate a normal distribution. For each cohort, the data were divided into five folds for cross-validation, with random splits repeated ten times. Given the weak linear discriminative capability of features extracted by the foundation model in this high-dimensional prediction task, a hidden layer with 512 dimensions was added to the penultimate layer on top of various WSI-based feature fusion operations. Considering the complexity of the task, a correlation loss term was added in addition to the MSE loss. The correlation loss computes the

Pearson correlation coefficient for each output dimension, averages them, and subtracts the result from 1, where a lower loss indicates higher correlation. To balance the smaller magnitude of the correlation loss, which ranges from 0 to 1, compared to the MSE in the high-dimensional prediction task, a weight of 50 was introduced. The batch size was all set to 256. For the BRCA and NSCLC cohort, the number of training epochs was set to 300, and the learning rate followed a cosine annealing schedule, decaying from 1e-4 to 1e-6 over a single cycle. For the CRC cohort, the number of training epochs was set to 800, while retaining the same learning rate schedule.

1. Oord, A.v.d., Y. Li, and O. Vinyals, *Representation learning with contrastive predictive coding.* arXiv preprint arXiv:1807.03748, 2018.
